# Glomerular Hyperfiltration, Charge Selectivity, and the Low-Dimensional Structure of Glomerular Transport

**DOI:** 10.64898/2026.06.23.733946

**Authors:** Carl M. Öberg

**Author notes:** **Correspondence:** Carl Öberg, Njurmedicin SUS Lund, Barngatan 2a, SE-221 85 LUND, Sweden.

## Abstract

**Background:** The relative contributions of molecular size, electrostatic charge, and filtration rate to glomerular transport remain controversial. We hypothesized that glomerular sieving data contain a limited number of underlying transport modes that can be identified directly from experimental measurements.

**Methods:** Glomerular sieving coefficients were measured in anesthetized rats using neutral and anionic polysucrose during baseline conditions and glucagon-induced hyperfiltration. Data were analyzed using aligned-rank two-factor ANOVA, nonlinear mixed-effects regression of an electrostatic distributed two-pore model, pairwise correlation analysis, and principal component analysis.

**Results:** Hyperfiltration reduced the sieving of small and intermediate polysucrose molecules, whereas anionic polysucrose exhibited lower sieving coefficients than neutral polysucrose over a broad range of molecular sizes. An electrostatic distributed two-pore model accurately reproduced the observed effects of filtration rate and molecular charge and yielded an effective pore-wall charge density of 5.4 mC/m² (95% confidence interval, 4.5–6.6). Pairwise correlation analysis revealed strong coupling between neighboring molecular sizes throughout the entire measured size range. Principal component analysis of the 2.5–8.0 nm size-selective region showed that the first principal component explained 96.3% of the variance and the first two principal components explained 99.9% of the variance. Separate analyses of the 2.5–5.0 nm and 5.0–8.0 nm transport regions showed that the first principal component explained 99.4% and 89.5% of the variance, respectively.

**Conclusions:** Glomerular sieving curves exhibited a highly constrained low-dimensional structure despite differences in molecular charge, filtration rate, and individual animals. The observed transport structure was consistent with distinct small-pore and large-pore transport domains and enabled highly effective principal component–based denoising of experimental sieving data.

**KEY POINTS:**

- Glomerular transport of low-molecular-weight polysucrose was almost perfectly one-dimensional.
- Two transport modes explained > 99% of variability across the complete glomerular sieving curve.
- Glomerular charge selectivity was consistent with electrostatic partitioning within a two-pore transport framework.

## INTRODUCTION

The glomerular filtration barrier (GFB) is a specialized three-layer structure that enables the kidney to selectively filter blood plasma. It consists of a fenestrated glomerular endothelium, the intervening glomerular basement membrane (GBM), and podocyte foot processes with their slit diaphragms ^1^. In concert, these layers ensure the free passage of water and small solutes while retaining most macromolecules like albumin ^2^. This selective filtration is governed largely by molecular size and shape, but it has long been postulated that electrostatic charge also plays a key role in hindering the passage of solutes.

Classic studies have supported the concept of a charge barrier in the GFB ^2,3^. Experiments with charged polysaccharide (dextran) polymers showed that negatively charged dextrans were restricted more than neutral ones of the same size, whereas cationic dextrans were filtered more readily ^4,5^. Charge selectivity has also been proposed as a key factor in explaining why native anionic proteins exhibit lower glomerular sieving coefficients (GSCs) compared to uncharged polysucrose (previously called Ficoll) molecules of similar size ^2,3^. In terms of GSC, 5.5 nm polysucrose approximately corresponds to 3.6 nm albumin ^6^. The electrokinetic model of glomerular filtration suggests that fluid flow through the charged filtration barrier generates a streaming potential proportional to the glomerular filtration rate, creating an electric field that repels negatively charged macromolecules like albumin ^7^. However, several studies have questioned whether the GFB’s charge-based filtration is as significant as once thought ^1,8^. The electrokinetic model has also been criticized ^8^. First, it relies on a reversed streaming potential – *i.e.* an electric field direction opposite to what classical theory would predict for a negatively charged capillary wall ^8^. Computational studies by Dechadilok & Deen suggest that electrokinetic effects play only a minor role in the hindrance of charged solute across charge membranes, with solute partitioning playing the key role ^9^.

Fundamental electrokinetic theory predicts that the magnitude of the electrokinetic potential across a charged membrane depends on the filtration rate ^10^. To test this physics-based prediction in the glomerular context, we measured glomerular permeability (sieving coefficients) for a neutral polysucrose and a similarly sized negatively charged (anionic) polysucrose under conditions of normal GFR and acutely elevated GFR. The glomerular sieving coefficients for polysucrose serve as a representative surrogate marker of glomerular permeability for plasma proteins such as albumin ^6,11^. This 2 × 2 experimental design (filtration rate: normal *vs.* high; solute charge: neutral *vs.* anionic) was analyzed with a two-way ANOVA to assess the main effects of filtration rate and solute charge, as well as their interaction. It should be noted that increasing the filtration rate is expected to decrease solute sieving also of neutral solutes ^2,6,12^.

## METHODS

### Animal Preparation

The experimental studies were performed in fifteen healthy male Wistar rats having an average body weight (BW) of 252 g (242-261g) with free access to food and water. All procedures were approved by the local Animal Ethics Committee at Lund University, Sweden. A total of 3 animals (number 7, 10 and 11) were excluded from the study due to sudden death, ureteral bleeding and blood sampling error. Anesthesia was induced by an intraperitoneal injection with pentobarbital (Pentobarbitalnatrium vet. APL 60 mg/ml), 1.5 mL/kg BW. Supplementary doses of pentobarbital (50-100 μL) were given via the tail artery as needed to maintain a surgical plane. A heating pad was used to maintain the body temperature at 37°C. A tracheotomy was performed (using a PE-240 tube) to facilitate breathing. The tail artery was cannulated (using a PE-50 cannula) and used for administration of horse serum, maintenance of anesthesia, and for continuous monitoring and registration of mean arterial blood pressure and heart rate (HR) on a data acquisition system (BioPack Systems model MP150 with AcqKnowledge, BioPack Systems Inc., Goleta, CA). Following a minimal abdominal incision, the left ureter was cannulated to allow direct urine collection. The left carotid artery was cannulated (PE-50) and used for blood sampling. The left and right internal jugular veins were cannulated (PE-50) for infusion of polysucrose and glucagon, respectively. All connectors, cannulas and tubes in contact with the circulation were prepared and flushed post administration with a dilute heparin (Heparin LEO 5000 IE/mL) solution (∼25 IE/mL in NaCl aq. 9 mg/mL) to prevent clotting. Following surgical preparation, the animals were allowed to stabilize for approximately 30 minutes. Adequacy of anesthesia and stability of hemodynamics (blood pressure and heart rate) were monitored throughout the experiment.

### Glomerular Filtration Markers

We used polysucrose as the test macromolecule to assess glomerular permeability. Polysucrose 70 kDa/400 kDa (TdB Consultancy, Uppsala, Sweden) was chosen for its wide range of molecular radii (∼1–8 nm) and electroneutral character. Carboxymethylation of neutral polysucrose was performed as described previously ^13^ resulting in anionic polysucrose. Previous work have shown that this anionic polysucrose is structurally the same as neutral polysucrose but carries a charge density comparable to that of albumin ^14,15^. Both neutral polysucrose and anionic polysucrose were fluorescently labeled with fluorescein isothiocyanate (FITC) to enable detection. Elugrams (Supplemental Figure 1) showed an elution/radius distribution very similar to that of native polysucrose, indicating that carboxymethylation did not cause any major detectable change in hydrodynamic size distribution.

### Experimental Protocol

Each rat underwent two experimental phases: a baseline filtration period at resting GFR, followed by a hyperfiltration period induced by volume expansion and glucagon ^12^. A 1:24 mixture of polysucrose-70 and polysucrose-400, or their anionic counterparts, was administered together with FITC-inulin as a bolus dose after surgery. Immediately thereafter, a continuous infusion of the same mixture FITC-polysucrose (neutral or anionic, depending on the experimental group) was begun via the jugular catheter at a rate of 50 μL/min to maintain a stable plasma polysucrose level. After allowing ∼20–30 minutes for equilibration of polysucrose between plasma and interstitial compartments, baseline urine and plasma samples were collected. Urine was collected via the ureteral catheter into pre-weighed tubes over a timed interval of 5 min, and an arterial blood sample was drawn at the midpoint (at 2.5 min) of the urine collection. Heparinized saline was flushed into the arterial line after each blood draw to prevent clotting. Glomerular filtration rate (GFR) during this baseline period was measured by the clearance of two independent filtration markers: FITC-inulin and radio-labeled chromium-51 EDTA. After the baseline collections, an intervention was performed to acutely raise the GFR (hyperfiltration phase). We implemented a volume expansion and hormonal stimulation protocol based on Rippe *et al.* ^12^. First, 5 mL of a colloid solution (sterile horse serum, Sigma-Aldrich) was infused intravenously over ∼10 minutes to expand plasma volume. Concurrently, a continuous infusion of glucagon (3 µg/min, Sigma) was started and maintained throughout the high-GFR period. Glucagon is known to dilate renal afferent arterioles and increase renal plasma flow, thereby augmenting GFR, while volume expansion raises cardiac output and renal perfusion pressure. Together, these maneuvers approximately double the GFR in rats. We continued the polysucrose (neutral or anionic) and inulin/EDTA infusions without interruption during this transition. After ∼15 minutes of glucagon+volume loading, a new steady state was achieved, and a second set of urine and plasma samples was collected (using the same durations and methods as in the baseline period). At the end of the experiment, the animal was euthanized with an overdose of potassium chloride.

### Analytical Methods

A high-performance size-exclusion chromatography (HPSEC) system (Waters, Milford, MA) was used to assess the concentrations of FITC-polysucrose and FITC-inulin in urine and plasma samples. An autosampler (Waters 717 Plus) loaded the samples onto the system, and the mobile phase was driven by a pump (Waters 1525). Fluorescence detection was performed using a fluorescence detector (Waters 2475) with an excitation wavelength of 492 nm and an emission wavelength of 518 nm. Size separation was achieved using an Ultrahydrogel 500 column (Waters), calibrated with polysucrose and protein standards as described previously ^16^. The sieving coefficients of FITC-polysucrose were determined as the fractional clearance (θ) using the following equation:

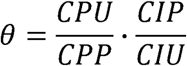

where CPP represents the concentration of polysucrose in plasma, CPU represents the polysucrose concentration in urine, CIP represents the inulin concentration in plasma, and CIU represents the inulin concentration in urine. Glomerular filtration rate (GFR) was measured in the left kidney using ^51^Cr-EDTA. GFR was calculated as the urinary excretion of ^51^Cr-EDTA and/or FITC-inulin per minute (Ut × Vu) divided by the plasma tracer concentration (Pt), where Ut represents the tracer concentration in urine and Vu is the urine flow rate per minute. A 25-μL blood sample was collected at the midpoint of each urine collection period to assess the tracer concentration in plasma. Hematocrit was measured throughout the experiment to convert blood radioactivity (^51^Cr-EDTA) to plasma radioactivity. Urine and blood radioactivities were quantified using a gamma counter (Wizard 1480, LKP, Wallac, Turku, Finland). Since the variability (coefficient of variation) of FITC-inulin-based GFR measurements was slightly higher than that of ^51^Cr-EDTA-based GFR measurements, we present GFR values obtained using ^51^Cr-EDTA throughout the study.

### Distributed Two-Pore Analysis

To provide a mechanistic interpretation of the observed sieving data, a distributed two-pore model of glomerular filtration was fitted to the experimental polysucrose sieving coefficients (for details, see Supplements). The model has previously been described in detail and represents the glomerular filtration barrier as a population of small pores with a log-normal size distribution and a small population of larger pores accounting for the filtration of the largest macromolecules ^17^. Electrostatic effects were modeled utilizing the semi-analytical solution by Smith and Deen ^18^. The model predicts the glomerular sieving coefficient as a function of molecular radius, glomerular filtration rate, hydraulic permeability, pore-size distribution and electrostatic interactions.

The small-pore radius was fixed at 3.66 nm, consistent with previous analyses of Ficoll transport in the rat glomerulus. Likewise, the geometric standard deviation of the small-pore population was fixed at 1.154 based on the highly consistent estimates obtained from preliminary analyses of individual animals. These parameters were therefore regarded as structural properties of the glomerular filtration barrier and are not expected to vary substantially between experimental groups.

Model fitting was performed using nonlinear mixed-effects regression (nlme package, R Foundation for Statistical Computing, Vienna, Austria). The hydraulic permeability parameter (*K_f_*) was allowed to vary between animals through a random intercept term, whereas the electrostatic pore-wall charge density parameter (*q_r_*) and the large-pore fraction parameter (*α_L_*) were treated as a fixed effect shared by all animals. Individual glomerular filtration rates measured by ^51^Cr-EDTA clearance were used as model inputs.

### Principal Component Analysis

To investigate the dimensional structure of glomerular transport, principal component analysis (PCA) was performed on log-transformed polysucrose sieving coefficients. Analyses were restricted to the size-selective region (2.5–8.0 nm), whereas the smallest polysucrose fraction (2.0 nm) was excluded because it approached free filtration and exhibited weaker coupling to the remaining size-selective transport domain.

The data matrix was organized with polysucrose radii as observations and individual experimental conditions (animals, charge states, and filtration states) as variables. Prior to PCA, sieving coefficients were transformed using the base-10 logarithm. Variables were centered using the column median and scaled using the interquartile range (IQR) to reduce sensitivity to outliers. Principal components were obtained using singular value decomposition.

The proportion of variance explained by each principal component was calculated from the corresponding singular values. To assess the dimensionality of glomerular transport, PCA was performed for the complete size-selective region (2.5–8.0 nm) as well as separately for the small-pore dominated (2.5–5.0 nm) and large-pore dominated (5.0–8.0 nm) regions.

For regularization, sieving curves were reconstructed using only the dominant principal component(s), thereby suppressing high-frequency fluctuations while preserving the major transport structure.

### Data and Statistical Analysis

All data are presented as median values with interquartile ranges (IQR) in parentheses. We employed a two-way analysis of variance (ANOVA) on aligned rank-transformed data (ART approach) to assess the effects of (i) Polysucrose charge (neutral vs anionic) and (ii) GFR condition (baseline vs high GFR) on the sieving coefficients. This non-parametric 2 × 2 ANOVA was implemented using the ARTool package in R (R Foundation for Statistical Computing, Vienna, Austria). Interaction between the two factors was included in the model. Based on our study design and sample size, we considered P ≤ 0.05 to indicate statistically significant main effects of charge or GFR. For the interaction term, a threshold of P ≤ 0.10 was used to denote a potential interactive trend, given the lower power to detect interaction effects. Post-hoc comparisons (if warranted) were planned using Wilcoxon signed-rank tests (for paired baseline vs high GFR comparisons within each polysucrose type) or Mann-Whitney U tests (for neutral vs anionic comparisons under a given condition), with Bonferroni adjustment for multiple comparisons. Statistical analyses were performed using R (version 4.3). All P values are two-tailed.

## RESULTS

### Baseline parameters

Glomerular sieving of neutral and anionic polysucrose was studied in healthy anesthetized rats during baseline conditions and during experimentally induced hyperfiltration. After a stabilization period of 20 min, renal filtration markers were measured during a control period and again following glucagon administration combined with volume loading to increase glomerular filtration rate. Six animals were studied in each group. Basic physiological parameters were comparable between groups. Body weight averaged 249 g (neutral group) and 256 g (anionic group). Mean arterial pressure remained stable between baseline and hyperfiltration periods (neutral: 99 *vs.* 102 mmHg; anionic: 106 *vs.* 108 mmHg). Heart rate increased during hyperfiltration in both groups (neutral: 324 *vs.* 395 bpm; anionic: 332 *vs.* 367 bpm), consistent with the systemic effects of glucagon.

### Effects on Glomerular Filtration Rate

Glomerular filtration rate increased markedly during glucagon and volume loading in both groups (Table 1). Baseline GFR was 0.55 (0.45–0.64) mL/min in rats receiving neutral polysucrose and 0.54 (0.48–0.62) mL/min in rats receiving anionic polysucrose. Following glucagon and volume loading, GFR increased to 0.91 (0.88–0.96) mL/min and 0.92 (0.90–1.03) mL/min, respectively. Similar changes were observed when GFR was estimated using FITC-inulin clearance. A nonparametric two-factor analysis (aligned rank transform) confirmed a strong effect of hyperfiltration on GFR (*F*_1,10_ = 74.0, *p* = 6.2 × 10^-6^), with no effect of polysucrose charge (*p* = 0.84) and non-significant charge × hyperfiltration interaction (*p* = 1.00). In summary, volume loading and glucagon strongly increased GFR, groups were hemodynamically similar and the hyperfiltration response was near identical between groups.

**Table 1.**
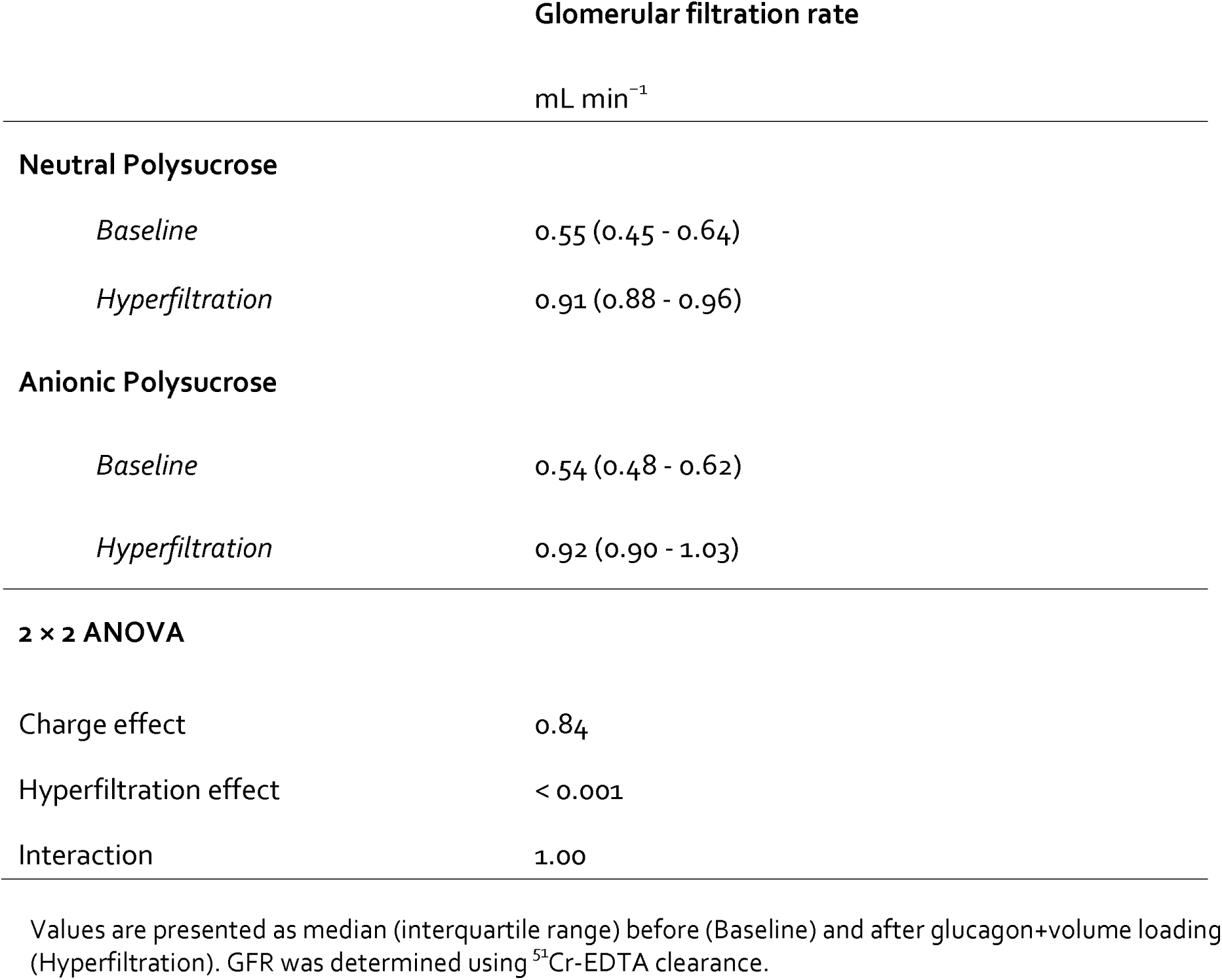
Glomerular filtration rate (mL/min) at baseline and during glucagon-induced hyperfiltration in rats receiving neutral or anionic polysucrose.

### Glomerular sieving of neutral and anionic polysucrose

Glomerular sieving coefficients (θ) for polysucrose molecules for three different hydrodynamic radii (2.5, 3.6 and 6.4 nm) are summarized in Table 2. At baseline, filtration was strongly size-dependent in both groups, with high sieving coefficients for smaller molecules and progressively lower values for larger molecules. For the 2.5 nm fraction, baseline sieving coefficients were 0.64 (IQR 0.60–0.66) in the neutral polysucrose group and 0.51 (IQR 0.50–0.52) in the anionic group. During glucagon-induced hyperfiltration, sieving coefficients decreased markedly in both groups to 0.44 (0.39–0.45) and 0.43 (0.41–0.46), respectively. Nonparametric factorial analysis confirmed a strong effect of hyperfiltration on the sieving of 2.5 nm molecules (*F*_1,10_ = 100.4, *p* = 1.6 × 10^-6^), together with a significant interaction between polysucrose charge and treatment condition (*F*_1,10_ = 41.4, *p* = 7.5 × 10^-5^) meaning that the effect of effect of charge is dependent on whether there is hyperfiltration or not.

**Table 2.**
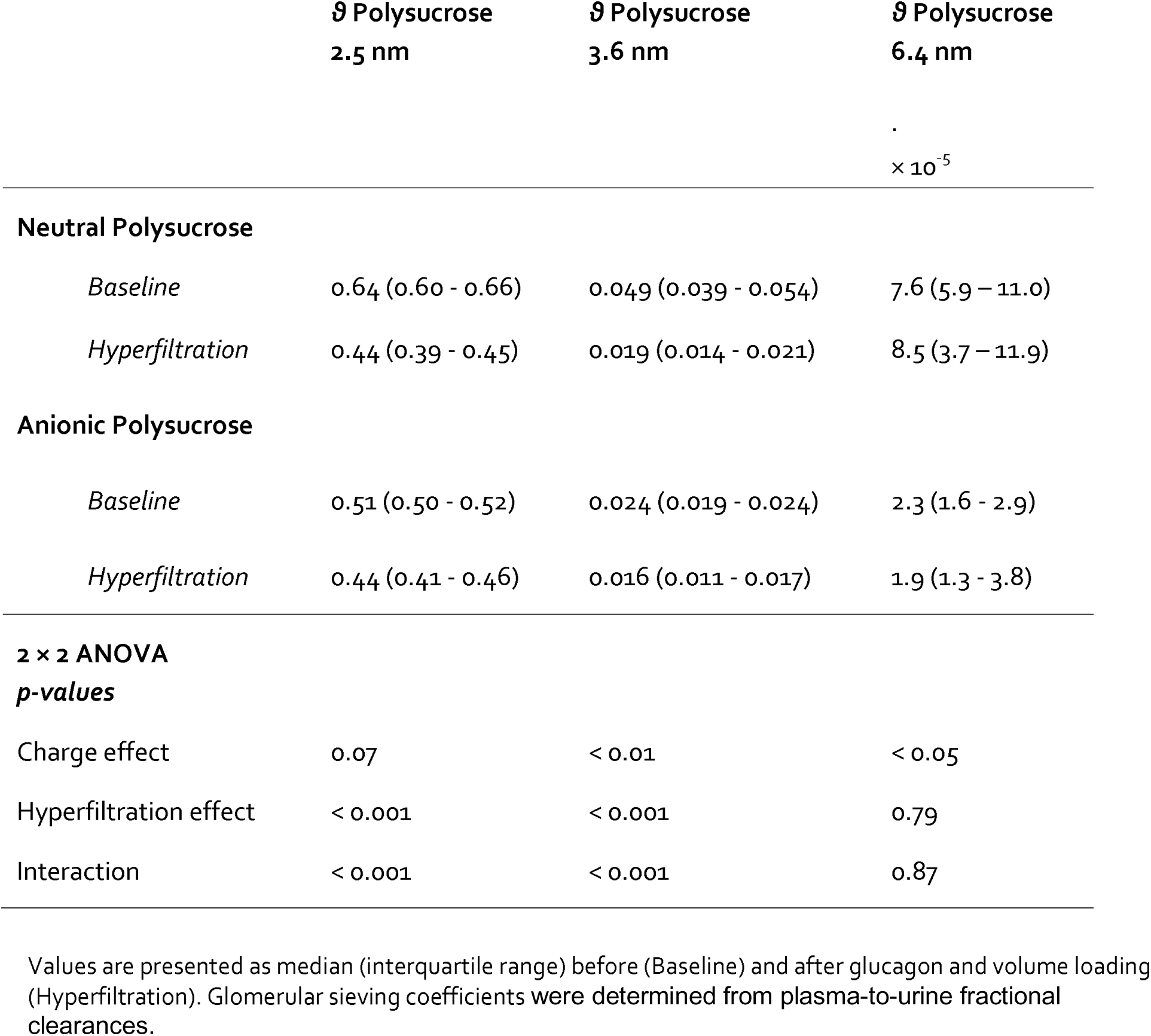
Glomerular sieving coefficients for small- (2.5 nm), intermediate- (3.6 nm) and large (6.4 nm) molecular polysucrose.

For the intermediate-sized molecules (3.6 nm), baseline sieving coefficients were lower and differed between groups, with values of 0.050 (0.039–0.055) for neutral polysucrose and 0.024 (0.019–0.024) for anionic polysucrose. Hyperfiltration reduced sieving coefficients in both groups to 0.019 (0.014–0.022) and 0.016 (0.012–0.017), respectively. Statistical analysis again demonstrated a significant effect of hyperfiltration (*F*_1,10_ = 128.0, *p* = 5.1 × 10⁻⁷), a significant interaction between charge and treatment (*F*_1,10_ = 30.5, *p* = 2.5 × 10⁻⁴), and an overall effect of charge (*F*_1,10_ = 17.3, *p* = 0.0019).

For the largest molecules studied (6.4 nm), sieving coefficients were expectedly low under all conditions, consistent with strong glomerular size restriction. Median values were 0.08 ×10^-3^ (0.06–0.11 ×10^-3^) for neutral polysucrose and 0.03 ×10^-3^ (0.01–0.03 ×10⁻³) for anionic polysucrose at baseline, and hyperfiltration did not significantly alter sieving for this molecular size (treatment effect *F*_1,10_ = 0.07, p = 0.79; interaction *F*_1,10_ = 0.03, p = 0.87), although a modest overall effect of charge was present (*F*_1,10_ = 6.64, p = 0.028).

### Mechanistic analysis of glomerular sieving

To investigate whether the observed effects of molecular size, charge and filtration rate could be explained within a common mechanistic framework, the complete polysucrose sieving curves were analyzed using an electrostatic distributed two-pore model. Model predictions obtained by nonlinear mixed-effects regression are shown in Figure 2.

**Figure 1.**
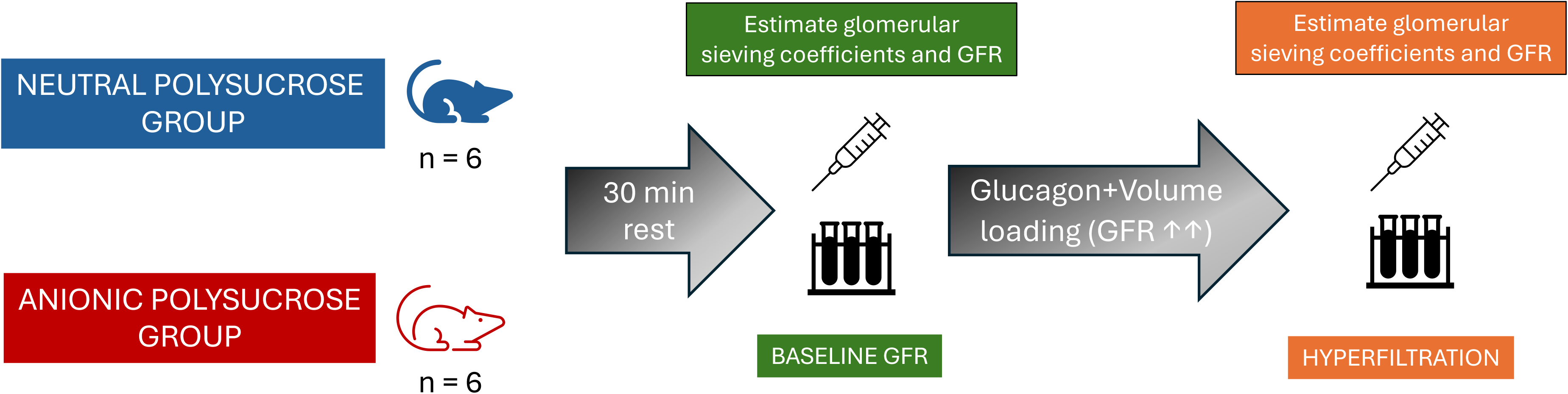
Schematic diagram of the experimental setup. Schematic of the experimental protocol in two groups of anesthetized rats receiving neutral or anionic polysucrose infusions. Baseline glomerular filtration rate (GFR) and sieving coefficients are measured, followed by a 30-min rest period. GFR is then increased via glucagon + volume loading, and measurements are repeated under hyperfiltration.

**Figure 2.**
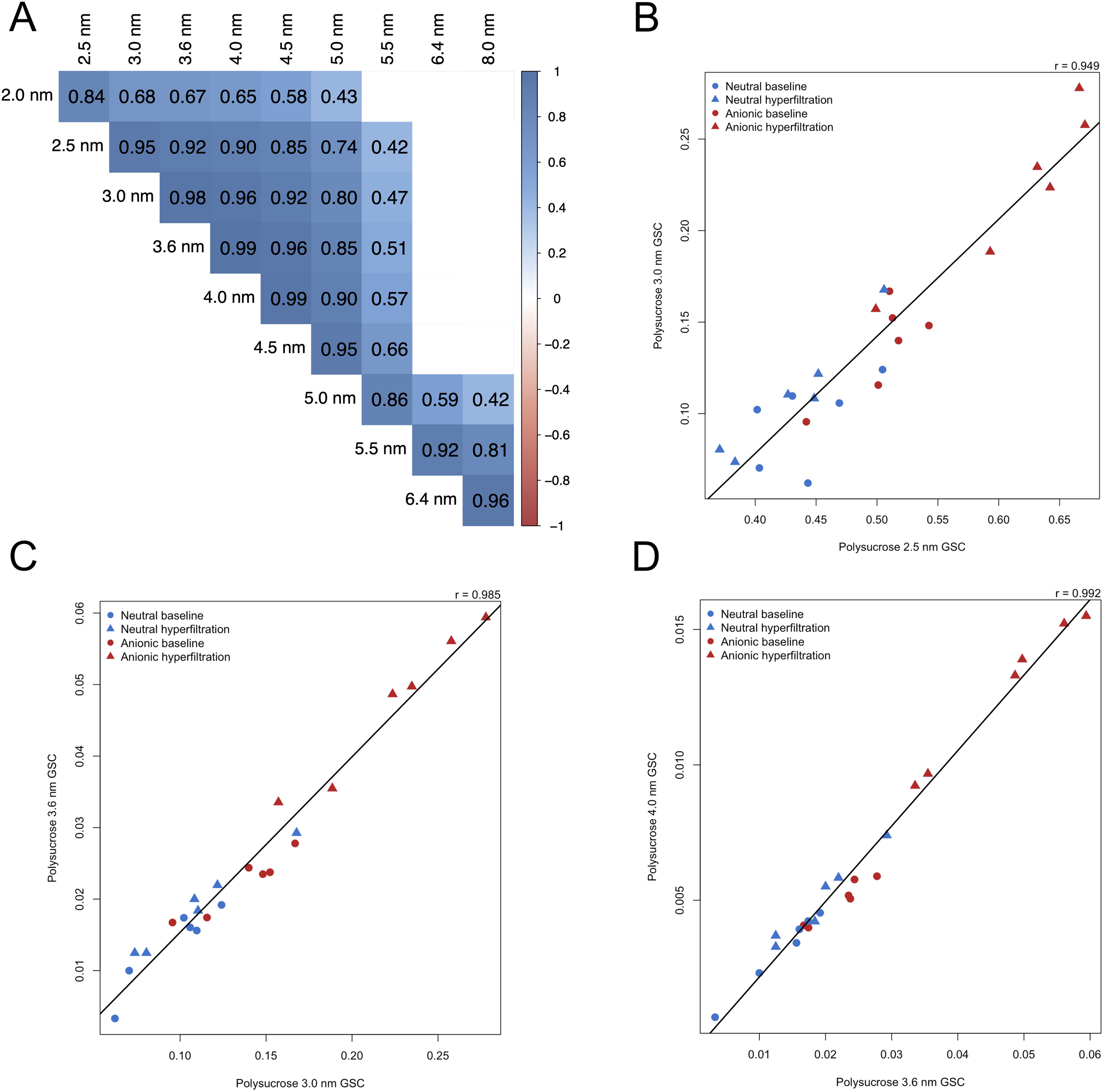
Glomerular sieving coefficients for neutral and anionic polysucrose during normofiltration and hyperfiltration. Glomerular sieving coefficients (θ) are shown as a function of hydrodynamic radius for neutral polysucrose (bottom panels) and anionic polysucrose (top panels) under baseline conditions (right panels) and glucagon-induced hyperfiltration (left panels). Grey circles represent individual animals (n=6 per group), blue circles represent group median values and red lines show predictions from the electrostatic distributed two-pore model obtained by nonlinear mixed-effects regression. The model accurately reproduced the observed size-dependent decline in sieving coefficients over approximately five orders of magnitude and captured the effects of both increased filtration rate and polysucrose charge. The largest polysucrose fractions (6.4–8.0 nm), which are dominated by the rare large-pore pathway, exhibited greater inter-animal variability than smaller molecules.

The model provided an excellent description of the experimental data across the full range of molecular radii studied. Predicted sieving coefficients closely followed the observed group medians for both neutral and anionic polysucrose under baseline conditions as well as during glucagon-induced hyperfiltration. In particular, the model accurately reproduced the reduction in sieving coefficients observed during hyperfiltration for intermediate-sized polysucrose molecules and the lower permeability of anionic relative to neutral polysucrose.

Agreement between model predictions and experimental observations was particularly strong throughout the small-pore dominated region of the sieving curve (approximately 2.0–5.5 nm). Greater variability was observed among the largest polysucrose fractions (6.4–8.0 nm), consistent with a larger error of measurement in this size range. Nevertheless, the model successfully captured the overall behavior of both neutral and charged polysucrose transport over approximately five orders of magnitude in sieving coefficients.

The nonlinear mixed-effects analysis yielded physiologically plausible parameter estimates (Table 3). The estimated hydraulic permeability parameter was *K*_f_ = 0.10 mL/min/mmHg (95% CI 0.067–0.149), the electrostatic pore-wall charge parameter was |*q*_r_| = 0.0054 C/m^2^ (95% CI 0.0045–0.0065), and the large-pore fraction parameter was a = 2.8·10^-5^ (95% CI 2.2·10^-5^ to 3.6·10^-5^). In conclusion, the same model structure was able to reproduce both the effects of increased filtration rate and the effects of molecular charge, suggesting that these phenomena can be understood within a common transport framework.

**Table 3.**
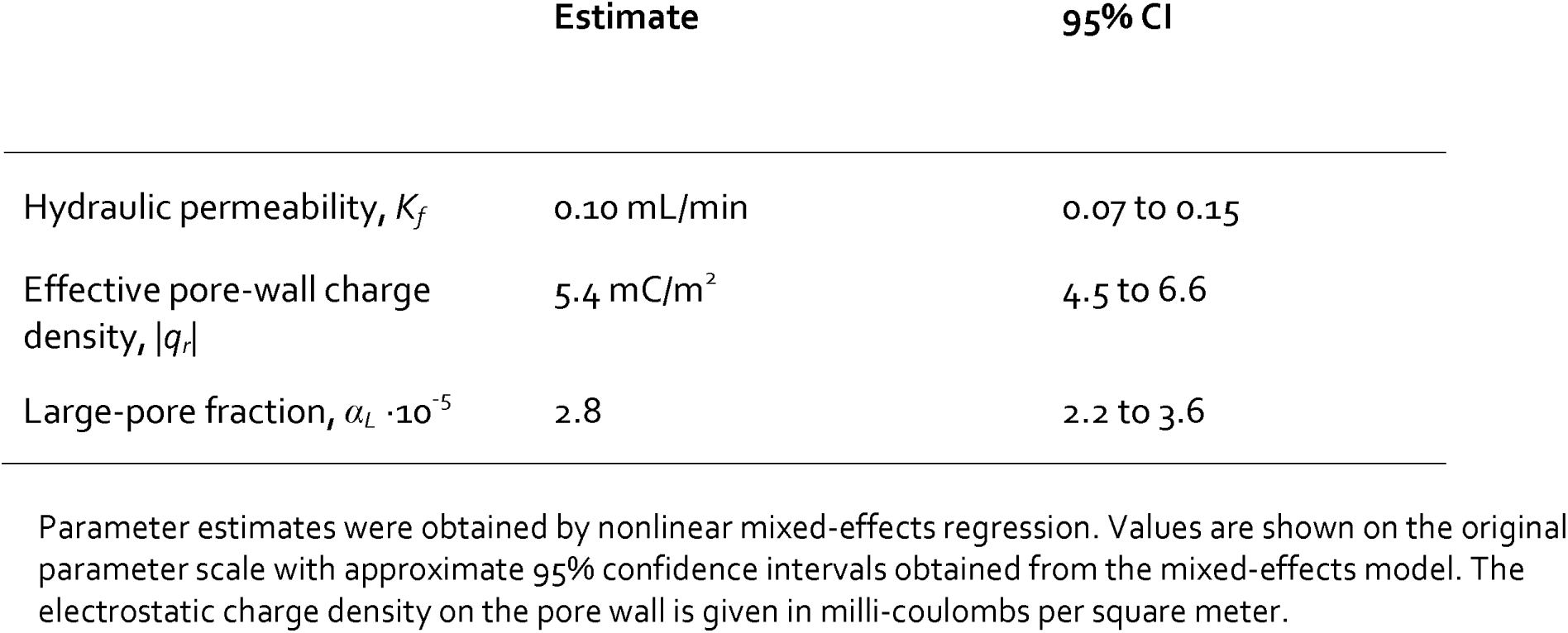
Electrostatic distributed two-pore model parameters.

### Correlation structure of glomerular sieving coefficients

Pairwise correlations between glomerular sieving coefficients revealed a remarkably structured pattern (Figure 3, Supplemental Figure 2). Strong positive correlations were observed between neighboring polysucrose fractions throughout the entire molecular size range. Within the small-pore dominated region (2.5–5.0 nm), correlation coefficients between adjacent molecular sizes typically exceeded 0.9 and frequently approached unity. Thus, knowledge of the sieving coefficient for a single polysucrose fraction largely determined the sieving coefficients of neighboring molecular sizes.

**Figure 3.**
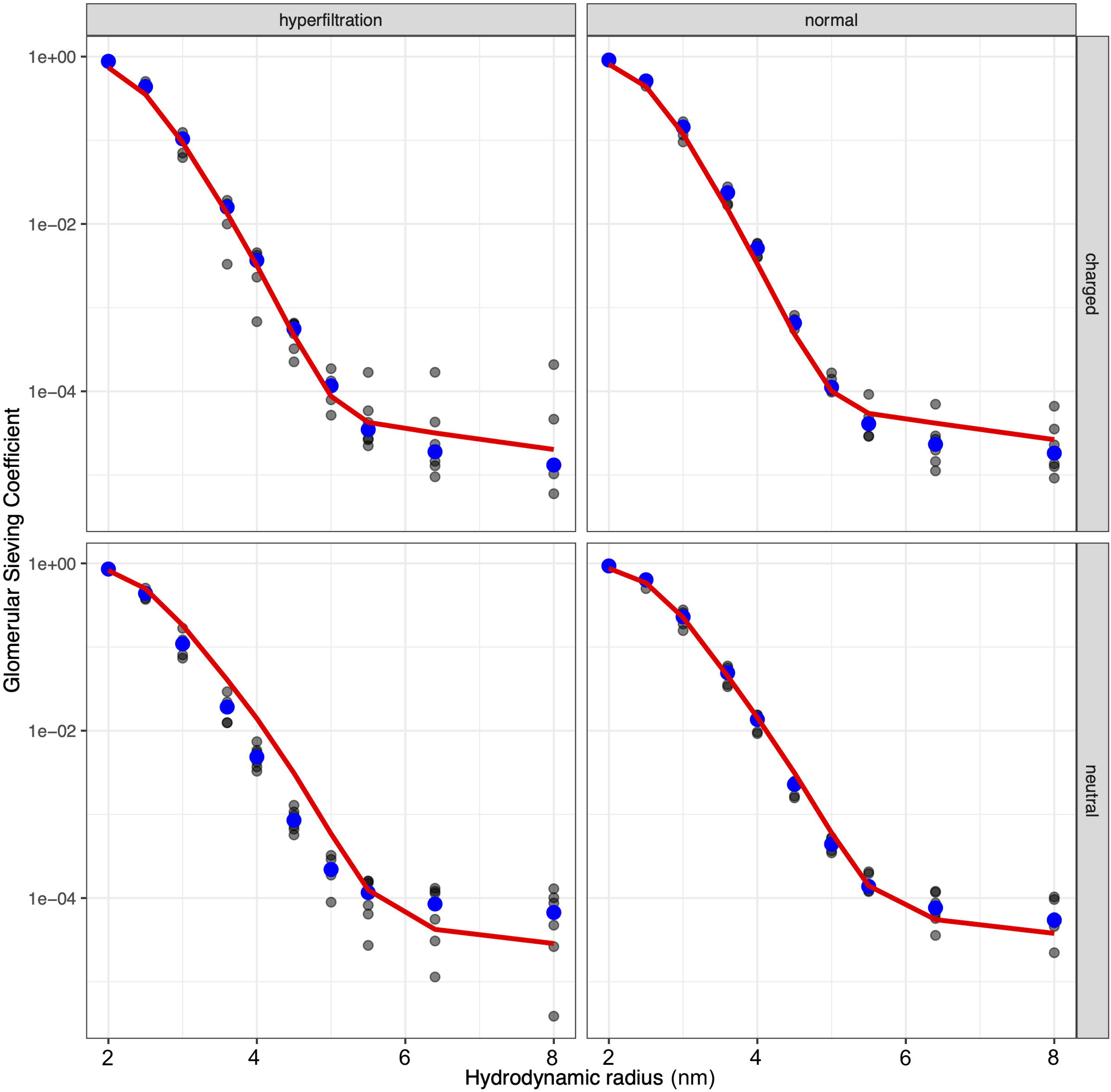
Low-dimensional structure of glomerular transport revealed by correlations between neighboring sieving coefficients. (A) Pairwise Pearson correlation coefficients between glomerular sieving coefficients measured for polysucrose fractions of different hydrodynamic radii. Strong correlations were observed throughout the small-pore dominated transport region (2.5–5.0 nm), whereas weaker correlations were observed between the small-pore and large-pore regions, consistent with the existence of two distinct transport domains. (B–D) Representative relationships between neighboring polysucrose fractions within the small-pore transport region. Each point represents an individual observation (n = 24), including neutral polysucrose and anionic polysucrose during both baseline filtration and glucagon-induced hyperfiltration. Blue symbols denote neutral polysucrose and red symbols denote anionic polysucrose; circles indicate baseline filtration and triangles indicate hyperfiltration. Solid lines represent least-squares linear regression. Despite substantial differences in molecular charge and glomerular filtration rate, neighboring sieving coefficients remained strongly coupled, with Pearson correlation coefficients ranging from 0.949 to 0.992. The preservation of these relationships across experimental conditions suggests that hyperfiltration and charge selectivity primarily move transport along a common low-dimensional transport manifold rather than generating distinct transport states.

Importantly, these relationships were observed despite combining neutral and anionic polysucrose, baseline and hyperfiltration conditions, and all individual animals in a single analysis. The strong correlations therefore cannot be explained by a single experimental condition but instead reflect an intrinsic property of glomerular transport.

The correlation matrix exhibited two highly correlated domains. A first domain encompassed small and intermediate polysucrose molecules (approximately 2.5–5.0 nm), whereas a second domain was observed among the largest polysucrose fractions (5.0–8.0 nm). Correlations were strongest within each domain and weaker between domains. This pattern suggests that transport of small and intermediate molecules is governed by a common underlying mechanism, whereas the largest polysucrose fractions contain additional information not captured by the small-molecule transport regime.

Collectively, these observations indicate that glomerular sieving curves occupy a substantially lower-dimensional space than would be expected if individual molecular sizes behaved independently. This hypothesis was explored further using principal component analysis.

### Principal component analysis of glomerular sieving curves

Principal component analysis was performed on the log-transformed polysucrose sieving curves over the size-selective range of 2.5–8.0 nm. This interval was chosen to exclude the smallest polysucrose fraction, which approached free filtration, and to focus the analysis on the region where size-selective restriction was most informative.

The PCA revealed a highly constrained structure of glomerular transport (Table 4). The first principal component accounted for 96.3% of the total variance, while the second component accounted for an additional 3.6%. Together, the first two components explained 99.9% of the variance, whereas all remaining components contributed only minimally. Thus, despite inclusion of neutral and anionic polysucrose, baseline and hyperfiltration conditions, and all individual animals, the complete glomerular sieving curve over the 2.5–8.0 nm range could be described almost entirely by two latent transport modes.

**Table 4.**
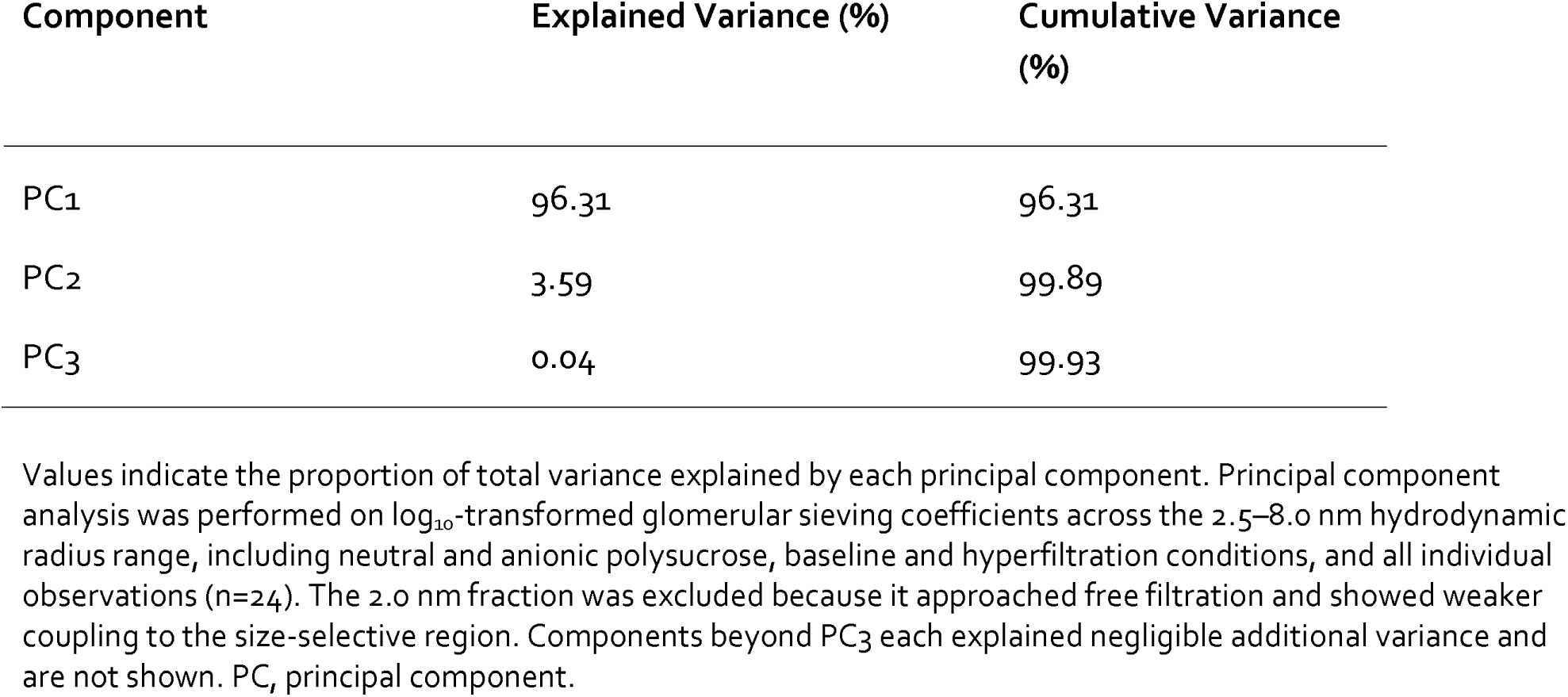
Principal component structure of glomerular polysucrose sieving coefficients in the size-selective range of 2.5–8.0 nm.

These findings demonstrate that glomerular sieving data occupy a remarkably low-dimensional space. Rather than behaving as independent measurements at each solute size, the sieving coefficients appear to be constrained by a small number of coordinated transport mechanisms.

Separate principal component analysis of the small-pore dominated 2.5–5 nm transport region revealed an almost perfectly one-dimensional structure (PC1 explained 99.4% of the total variance). Expectedly, projection onto the first principal component resulted in nearly identical curves compared to measured data (Supplemental Figure 3). Separate analysis for the large-pore dominated region (5–8 nm) showed that the first principal component accounted for 89.5% of the total variance. Although lower than observed for small solutes, this result still indicates a highly constrained transport structure. The residual variance was distributed across several minor components and largely reflected the lower signal-to-noise ratio associated with very low sieving coefficients in this size range. Projection onto the first principal component acted as a highly effective denoising procedure (Supplemental Figure 4).

## DISCUSSION

### The present study provides three major findings

First, glomerular hyperfiltration reduced the sieving of small and intermediate polysucrose molecules irrespective of solute charge. This finding is consistent with classical pore theory and previous Ficoll studies demonstrating decreased sieving coefficients at higher filtration rates ^6,12,19^. Importantly, the effect was observed for both neutral and anionic polysucrose, indicating that changes in glomerular filtration rate and electrostatic charge do not represent competing mechanisms but rather act within the same transport framework.

Second, negatively charged polysucrose exhibited significantly lower sieving coefficients than neutral polysucrose over a broad range of molecular sizes. The effect persisted during hyperfiltration and was accurately reproduced by an electrostatic distributed two-pore model. Because electrokinetic forces are predicted to scale with filtration rate, the absence of a marked GFR-dependent amplification of charge selectivity argues against a dominant role for streaming-potential effects. Instead, the data are more consistent with electrostatic partitioning at the plasma–barrier interface. This interpretation agrees with theoretical analyses by Dechadilok and Deen showing that electrostatic partitioning dominates electrokinetic contributions under physiologically relevant conditions ^9^.

Third, glomerular sieving curves exhibited a remarkably low-dimensional structure. Pairwise correlations between neighboring polysucrose fractions remained strong despite combining neutral and anionic polysucrose, baseline and hyperfiltration conditions, and all individual animals. Principal component analysis revealed that a single component explained 99.4% of the variance within the 2.5–5.0 nm size-selective region, whereas two components explained virtually all variance across the complete 2.5–8.0 nm range. Thus, glomerular transport appears to be governed by a very small number of coordinated transport modes rather than a large number of independent mechanisms.

The observed PCA structure is particularly noteworthy because it emerged directly from the experimental data before any pore-model assumptions were imposed. The correlation structure itself suggested the existence of two transport domains corresponding broadly to the classical small-pore and large-pore pathways ^17^. Principal component analysis subsequently quantified this observation. Within each transport domain, the dominant mode was effectively one-dimensional, implying the existence of implying the existence of a low-dimensional family of sieving curves from which most observed transport behavior can be reconstructed.

From a methodological perspective, the low-dimensional structure enabled highly effective PCA-based regularization of experimental sieving curves. Projection onto the dominant principal component substantially reduced high-frequency fluctuations while preserving the underlying transport curve. Such regularization may improve the stability of inverse pore-model analyses and could prove useful in future studies involving noisy macromolecular transport data.

Several limitations should be acknowledged. The study was performed in healthy rats under acute experimental conditions and may not fully reflect chronic hyperfiltration states or human glomerular disease. Furthermore, the principal component analysis was restricted to the size-selective region of the sieving curve. Nevertheless, the consistency of the findings across charge states, filtration rates and individual animals suggests that the observed low-dimensional transport structure reflects a fundamental property of glomerular filtration. In conclusion, the preservation of pairwise relationships between neighboring sieving coefficients across changes in filtration rate and solute charge suggests that these perturbations move transport along a common low-dimensional manifold rather than creating distinct transport states.

### Data sharing statement

All original data reported in this article have been deposited in Dryad and can be accessed online via http://doi.org/10.5061/dryad.wh70rxx45.

## Acknowledgements

Anna Rippe is gratefully acknowledged for excellent experimental work and HPLC-analyses of polysucrose.

## Supplemental material

This article contains the following supplemental material online:

**Supplemental Table 1.** Glomerular filtration rates (mL/min) at baseline and during glucagon-induced hyperfiltration in rats receiving neutral or anionic polysucrose

**Supplemental Table 2.** Glomerular sieving coefficients (θ) for neutral and anionic polysucrose molecules of different hydrodynamic radii during baseline conditions and glucagon-induced hyperfiltration.

**Supplementary Figure 1.** Pairwise relationships between glomerular sieving coefficients reveal a two-domain transport structure.

**Supplementary Figure 2.** Pairwise relationships between glomerular sieving coefficients reveal a two-domain transport structure.

**Supplementary Figure 3.** PCA-based regularization of glomerular sieving curves in the small-pore transport region (2.5–5.0 nm).

**Supplementary Figure 4.** PCA-based regularization of glomerular sieving curves in the large-pore transport region (5.0–8.0 nm).

**Electrostatic Distributed Two-pore Model**

