## Supplemental Material for "Glomerular Hyperfiltration, Charge Selectivity, and the Low-Dimensional Structure of Glomerular Transport"

Carl Öberg [1] M.D. Ph.D.

[1] Nephrology Division, Department of Clinical Sciences Lund, Skåne University Hospital, Lund, Sweden.


| CONTENTS | PAGE |
|---|---|
| **Supplemental Table 1.** Glomerular filtration rate (mL/min) at baseline and during glucagon-induced hyperfiltration | 2 |
| **Supplemental Table 2.** Glomerular sieving coefficients (θ) for neutral and anionic polysucrose molecules of different hydrodynamic radii during baseline conditions (normofiltration) and glucagon-induced hyperfiltration. | 3 |
| **Supplemental Figure 1.** Hydrodynamic size distributions of neutral and charge-modified FITC-polysucrose. | 4 |
| **Supplementary Figure 2.** Pairwise relationships between glomerular sieving coefficients reveal a two-domain transport structure. | 5 |
| **Supplementary Figure 3.** PCA-based regularization of glomerular sieving curves in the small-pore transport region (2.5–5.0 nm). | 7 |
| **Supplementary Figure 4.** PCA-based regularization of glomerular sieving curves in the large-pore transport region (5.0–8.0 nm). | 10 |
| **Electrostatic Distributed Two-pore Model** | 12 |
| **References** | 16 |

**Supplemental Table 1.** Glomerular filtration rate (mL/min) at baseline and during glucagon-induced hyperfiltration in rats receiving neutral or anionic polysucrose

| | **Neutral Polysucrose** *Baseline* | **Neutral Polysucrose** *Hyperfiltration* | | **Anionic Polysucrose** *Baseline* | **Anionic Polysucrose** *Hyperfiltration* |
|---|---|---|---|---|---|
| 1 | 0.53 | 0.90 | 1 | 0.59 | 1.07 |
| 2 | 0.42 | 1.08 | 2 | 0.63 | 0.78 |
| 3 | 0.66 | 0.97 | 3 | 0.48 | 0.91 |
| 4 | 0.40 | 0.87 | 4 | 0.74 | 1.07 |
| 5 | 0.70 | 0.91 | 5 | 0.43 | 0.92 |
| 6 | 0.58 | 0.78 | 6 | 0.49 | 0.89 |
| | **0.55 (0.45 - 0.64)** | **0.91 (0.88 - 0.96)** | | **0.54 (0.48 - 0.62)** | **0.92 (0.90 - 1.03)** |

Values are presented as individual animals with group median (interquartile range) shown in the final row (n = 6 per group) at before (Baseline) and after glucagon+volume loading (Hyperfiltration). GFR was determined using $^{51}Cr$-EDTA clearance.

**Supplemental Table 2.** Glomerular sieving coefficients (θ) for neutral and anionic polysucrose molecules of different hydrodynamic radii during baseline conditions (normofiltration) and glucagon-induced hyperfiltration.

| | | ***Neutral polysucrose*** | | | | **Anionic polysucrose** | | |
|---|---|---|---|---|---|---|---|---|
| | | *θ* *2.5 nm* | *θ* *3.6 nm* | *θ x 10⁻³* *6.4 nm* | | *θ* *2.5 nm* | *θ* *3.6 nm* | *θ* *6.4 nm* |
| **Normofiltration** | 1 | 0.63 | 0.050 | 0.12 | 1 | 0.51 | 0.024 | 0.12 |
| | 2 | 0.50 | 0.034 | 0.06 | 2 | 0.44 | 0.017 | 0.06 |
| | 3 | 0.59 | 0.035 | 0.09 | 3 | 0.54 | 0.023 | 0.09 |
| | 4 | 0.67 | 0.059 | 0.12 | 4 | 0.52 | 0.024 | 0.12 |
| | 5 | 0.67 | 0.056 | 0.04 | 5 | 0.50 | 0.017 | 0.04 |
| | 6 | 0.64 | 0.049 | 0.07 | 6 | 0.51 | 0.028 | 0.07 |
| | $\bar{x}$ | **0.64** **(0.60 - 0.66)** | **0.050** **(0.039 - 0.055)** | **0.08** **(0.06 - 0.11)** | $\bar{x}$ | **0.51** **(0.50 - 0.52)** | **0.024** **(0.019 - 0.024)** | **0.03** **(0.01 – 0.03)** |
| **Hyperfiltration** | 1 | 0.37 | 0.012 | 0.13 | 1 | 0.43 | 0.016 | 0.01 |
| | 2 | 0.38 | 0.012 | 0.06 | 2 | 0.40 | 0.010 | 0.01 |
| | 3 | 0.43 | 0.018 | 0.01 | 3 | 0.50 | 0.019 | 0.01 |
| | 4 | 0.51 | 0.029 | 0.11 | 4 | 0.47 | 0.016 | 0.04 |
| | 5 | 0.45 | 0.022 | 0.12 | 5 | 0.44 | 0.003 | 0.17 |
| | 6 | 0.45 | 0.020 | 0.03 | 6 | 0.40 | 0.017 | 0.02 |
| | $\bar{x}$ | **0.44** **(0.39 - 0.45)** | **0.019** **(0.014 - 0.022)** | **0.08** **(0.04 - 0.12)** | $\bar{x}$ | **0.43** **(0.41 - 0.46)** | **0.016** **(0.012 - 0.017)** | **0.01** **(0.01 - 0.03)** |

Values are presented as individual animals with group median (interquartile range) shown in the final row (n = 6 per group).

**Supplementary Figure 1: Hydrodynamic size distributions of neutral and charge-modified FITC-polysucrose.**

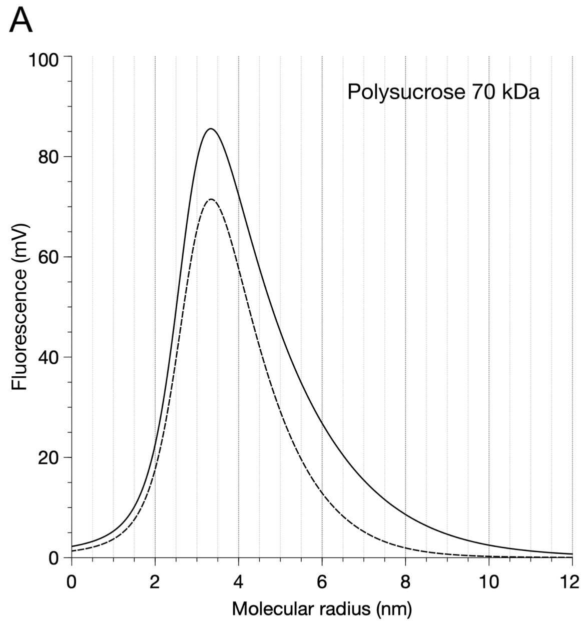


High-performance size-exclusion chromatography (HPSEC) elugrams are shown for FITC-labeled polysucrose 70 kDa (A) and FITC-labeled polysucrose 400 kDa (B). Solid lines represent neutral polysucrose and dashed lines represent charge-modified (anionic) polysucrose following carboxymethylation as described in the Methods section. The x-axis denotes the hydrodynamic radius corresponding to each elution volume following calibration with molecular size standards. The overall elution profiles were similar for neutral and charge-modified polysucrose, indicating that carboxymethylation produced substantial changes in molecular charge while preserving the overall hydrodynamic size distribution. These findings support the use of charge-modified polysucrose as a probe for electrostatic effects on glomerular transport without major concomitant alterations in molecular size.

**Supplementary Figure 2. Pairwise relationships between glomerular sieving coefficients reveal a two-domain transport structure.**

***Please note: Because of the large number of panels, the figure is best viewed electronically with magnification to appreciate the detailed correlation structure.***

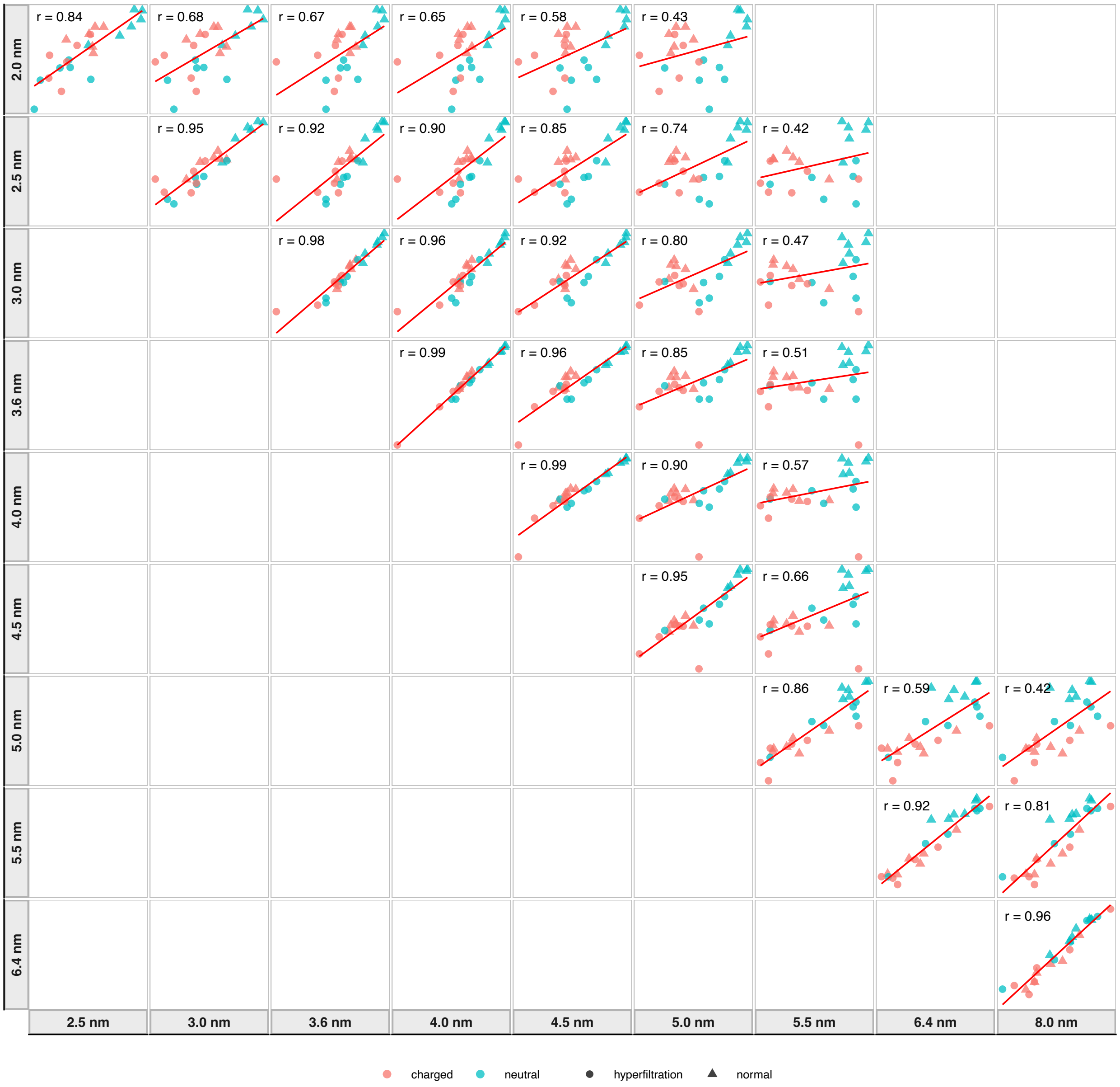


Each panel shows the relationship between two polysucrose sieving coefficients, including both neutral and anionic polysucrose during baseline conditions and glucagon-induced hyperfiltration (n=24 observations). For clarity, only a representative subset of polysucrose radii used in the pore-model analysis is shown. Points represent individual observations (red=charged, blue=neutral, triangles=hyperfiltration, circles=baseline) and solid lines indicate linear least-squares fits. Only statistically significant pairwise relationships are shown.

Strong correlations were observed between neighboring molecular sizes throughout the entire measured radius range, demonstrating that glomerular sieving coefficients do not behave as independent variables. Instead, knowledge of the sieving coefficient for one solute largely determines the sieving coefficients of neighboring solutes, despite substantial variation in molecular charge, filtration rate and individual animals.

The correlation structure furthermore revealed two distinct transport domains. Small and intermediate polysucrose fractions (approximately 2.0–5.0 nm) formed one highly correlated block, whereas the largest polysucrose fractions (5.5–8.0 nm) formed a second highly correlated block. Correlations were strongest within each block and markedly weaker between blocks. This pattern is consistent with the existence of two dominant transport pathways in the glomerular filtration barrier: a small-pore pathway governing transport of low- and intermediate-molecular-weight solutes and a large-pore pathway governing transport of the largest polysucrose fractions.

**Supplementary Figure 3. PCA-based regularization of glomerular sieving curves in the small-pore transport region (2.5–5.0 nm).**

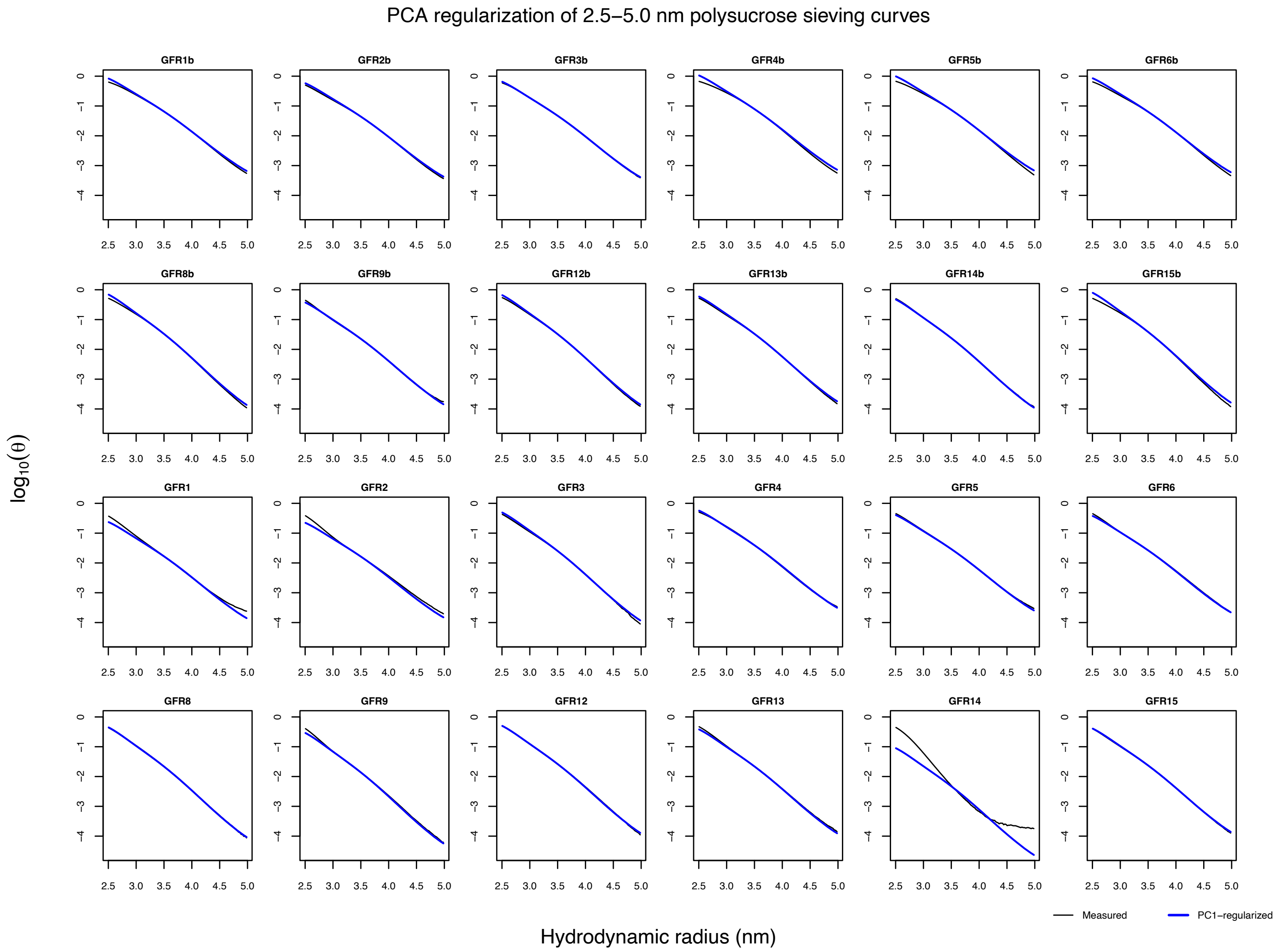

***Please note: Because of the large number of panels, the figure is best viewed electronically with magnification to appreciate the detailed correlation structure.***

Black curves represent measured log-transformed polysucrose sieving coefficients for each individual observation, including neutral and anionic polysucrose during baseline conditions and glucagon-induced hyperfiltration. Blue curves represent reconstruction using only the first principal component. The first and second rows represent neutral and anionic polysucrose during baseline conditions. The third and fourth row represent neutral and anionic polysucrose during hyperfiltration.

Principal component analysis of the complete 2.5–5.0 nm transport region revealed an almost perfectly one-dimensional transport structure, with the first principal component explaining 99.4% of the total variance. Consequently, reconstruction using a single component was sufficient to reproduce nearly all observed transport behavior despite substantial differences in molecular charge, filtration rate and individual animals. The close agreement between measured and reconstructed curves indicates that most variability in this size range can be described by a single underlying transport mode.

**Supplementary Figure 4. PCA-based regularization of glomerular sieving curves in the large-pore transport region (5.0–8.0 nm).**

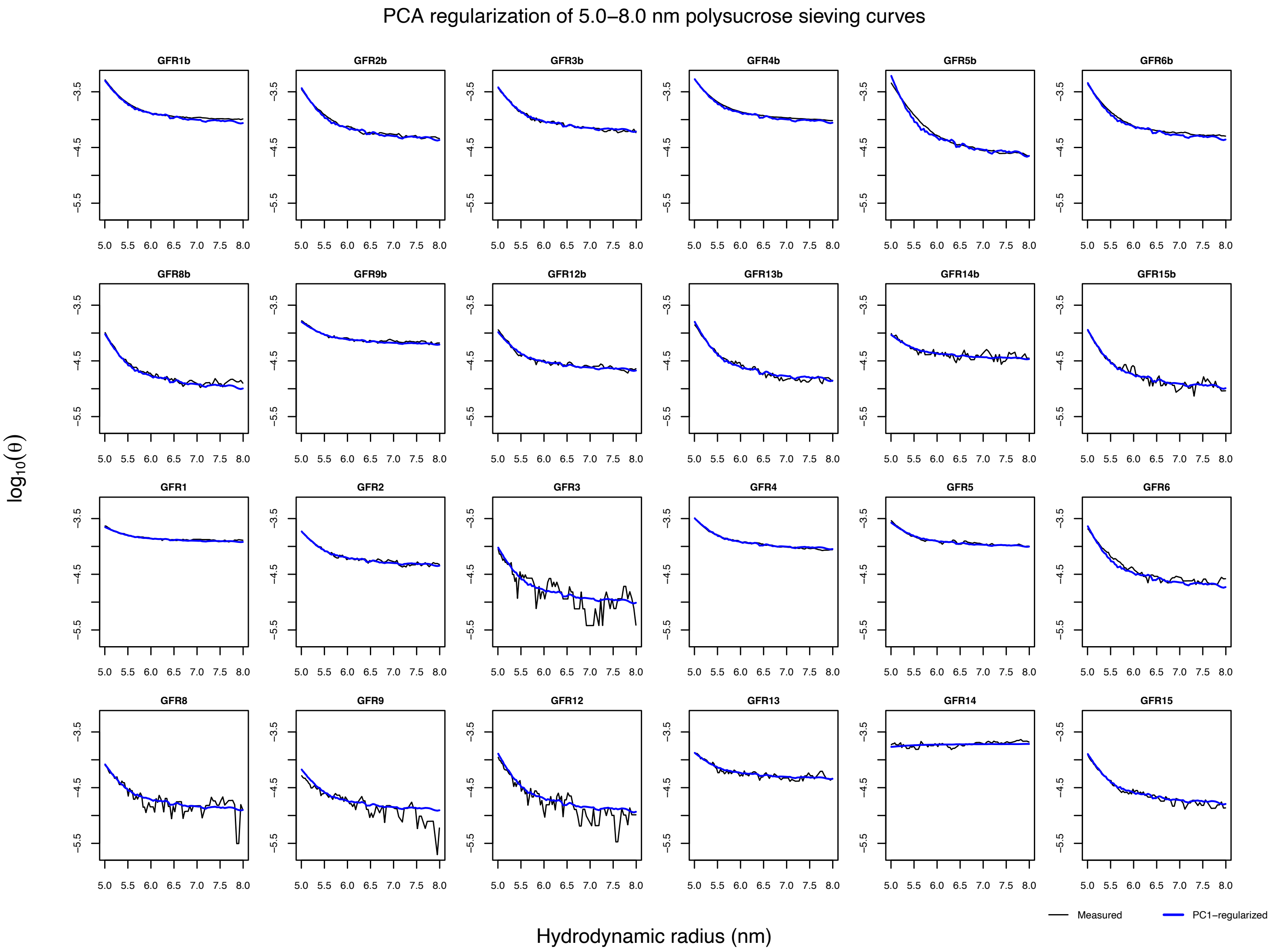

***Please note: Because of the large number of panels, the figure is best viewed electronically with magnification to appreciate the detailed correlation structure.***

Black curves represent measured log-transformed polysucrose sieving coefficients for each individual observation, including neutral and anionic polysucrose during baseline conditions and glucagon-induced hyperfiltration. Blue curves represent reconstruction using only the first principal component. The first and second rows represent neutral and anionic polysucrose during baseline conditions. The third and fourth row represent neutral and anionic polysucrose during hyperfiltration.

Although the first principal component accounted for a smaller fraction of the total variance in the large-pore region (89.5%) than in the small-pore region, reconstruction using a single component remained remarkably accurate. The principal effect of PCA regularization was suppression of high-frequency fluctuations while preserving the underlying transport curve. The residual variance was distributed across several minor components and is likely influenced by the substantially lower signal-to-noise ratio associated with very low sieving coefficients in this size range. One observation exhibited greater deviation from the common transport pattern than the remaining curves, but the overall agreement remained strong, supporting the existence of a highly constrained underlying large-pore transport structure.

### Electrostatic Distributed Two-pore Model

*Overview*

Glomerular sieving coefficients were analyzed using a distributed two-pore model extended to include electrostatic interactions between charged polysucrose molecules and the pore wall. The model builds on the distributed two-pore framework previously described for polysucrose transport, in which water and solute transport occur through a log-normally distributed small-pore pathway and a low-abundance large-pore pathway [1]. Electrostatic effects were incorporated by modifying the diffusive [2] and convective [3] restriction factors according to the electrostatic pore model of Smith and Deen [4].

For a solute with hydrodynamic radius *a*, the total glomerular sieving coefficient was calculated as the volume-flux weighted sum of the small- and large-pore contributions,

$$\theta(a) = (1 - f_L)\theta_S(a) + f_L\theta_L(a),$$

where $f_L$ is the fraction of filtrate passing through the large-pore pathway. For each pore pathway *p*, the sieving coefficient was calculated from the steady-state convection–diffusion equation,

$$\theta_p(a) = \frac{1 - \sigma_p(a)}{1 - \sigma_p(a)e^{-Pe_p(a)}},$$

where $\sigma_{p(a)}$ is the reflection coefficient and $Pe_{p(a)}$ is the Péclet number. The Péclet number was defined as

$$Pe_p(a) = \frac{J_{v,p}\left(1 - \sigma_p(a)\right)}{PS_p(a)},$$

where $J_{v,p}$ is the volume flux through pathway *p* and $PS_{p(a)}$ is the diffusive permeability–surface area product.

*Pore-size distribution*

The small-pore population was represented by a log-normal distribution of pore radii with geometric mean radius $r_S$ and geometric standard deviation $s_S$. In the main analysis, $r_S$ was fixed at 36.6 Å and $s_S$ at 1.154, based on previous rat polysucrose data [1] and preliminary fits showing highly consistent values across experimental conditions. The large-pore pathway was represented by a single pore radius $r_L$, fixed at 15 nm, and a large-pore flow fraction parameter $\alpha_L$.

For a log-normal pore-size distribution, moments of the pore radius distribution were evaluated analytically as

$$G_n = r_S^n e^{\frac{n^2(\log s_S)^2}{2}}.$$

These moments were used to calculate area- and flow-weighted quantities for the distributed small-pore pathway.

*Hydrodynamic hindrance factors*

In the absence of electrostatic interactions, diffusive and convective restriction were described using standard hydrodynamic hindrance factors as functions of

$$\lambda = \frac{a}{r_p},$$

where $r_p$ is the pore radius. The diffusive hindrance factor $H(\lambda)$ and convective hindrance factor $W(\lambda)=1-\sigma(\lambda)$ were set to zero when $\lambda \geq 1$. For $\lambda < 1$, polynomial approximations to the cylindrical pore hindrance functions were used [5].

For distributed pores, the effective diffusive restriction factor was calculated by averaging the single-pore diffusive hindrance over the pore distribution weighted by pore cross-sectional area. The effective convective restriction factor was calculated by averaging the single-pore convective hindrance weighted by hydraulic conductance, as previously described [1].

*Electrostatic interactions*

For anionic polysucrose, electrostatic interactions between the charged solute and charged pore wall were incorporated using the electrostatic framework of Smith and Deen [4]. The interaction was described by a solute surface charge density $q_s$, a pore-wall charge density parameter $q_r$, and the Debye length $\kappa^{-1}$. The Debye length was set to 0.79 nm, corresponding approximately to physiological ionic strength.

The surface charge density of anionic polysucrose was fixed at 15 mC/m$^2$. This value was used as an approximate effective charge density based on previous physicochemical estimates and the Grahame equation. Because the absolute value of $q_s$ is uncertain, the fitted pore-wall charge parameter $q_r$ should be interpreted as an effective electrostatic parameter rather than a direct physical measurement of fixed charge density. Changing the assumed value of $q_s$ would be expected to systematically rescale the fitted value of $q_r$, without materially changing the ability of the model to describe the observed sieving curves.

Electrostatic effects modified: 1. solute partitioning at the blood plasma/pore interface and 2. hindrance inside the pore. In general, the effects of electrostatic interactions on solute partitioning will outweigh that of intra-pore hindrance [2,6]. The electrostatic free energy $E(\beta)$, where $\beta$ is relative the radial position of the solute center within the accessible pore cross-section, was calculated using the Smith–Deen approximation. The partition coefficient was evaluated as

$$\Phi(a, r_p) = 2\int_0^{1-\lambda} \beta e^{\frac{-E(\beta)}{kT}}\, \mathrm{d}\beta,$$

where $E(\beta)/kT$ is the electrostatic potential energy per particle of the interaction between the solute and pore wall. More generally, the Boltzmann factor $E$ is a measure of the probability of finding a solute center at the radial position $\beta$.

The electrostatically modified diffusive hindrance factor was calculated as

$$H_z(a, r_p) = 2\int_0^{1-\lambda} K^{-1}\beta e^{\frac{-E(\beta)}{kT}}\,\mathrm{d}\beta,$$

Where $K^{-1}$ represents the reduced diffusional mobility (enhanced drag) inside the pore compared to that in bulk solution. The centerline approximation was used, which is a particularly good approximation in a charged pore [6]. The electrostatically modified convective hindrance factor [3] was calculated from

$$W_z(a, r_p) = 2\int_0^{1-\lambda} \beta(1-\beta^2) e^{\frac{-E(\beta)}{kT}}\,\mathrm{d}\beta.$$

For neutral polysucrose, $q_s = 0$, and the electrostatic terms reduce to the corresponding neutral hydrodynamic hindrance factors [5].

*Calculation of the electrostatic interaction energy*

The electrostatic interaction energy $E(\beta)$ between a charged spherical solute and a charged cylindrical pore wall was calculated according to the approximation of Smith and Deen [4]. Here, $\beta$ denotes the radial position of the solute center normalized to the pore radius. The dimensional interaction energy was written as

$$E(\beta) = r_p \varepsilon \left(\frac{kT}{e}\right)^2 \Delta G(\beta)$$

where $r_p$ is the pore radius, $\varepsilon$ is the permittivity of the medium, $k$ is Boltzmann's constant, $T$ is absolute temperature (310ºK), $e$ is the elementary charge and $\Delta G(\beta)$ is the dimensionless free-energy difference between the sphere–cylinder system and the isolated sphere and cylinder at infinite separation.

The dimensionless energy $\Delta G(\beta)$ was calculated as the sum of three terms representing solute self-energy, solute–pore interaction energy and pore-wall self-energy contributions, divided by the corresponding normalization factor, following Eq. 29 in Smith and Deen [4]. The auxiliary function $\Gamma(x)=\coth(x)-1/x$ was evaluated directly. The function $\Lambda(\beta, \tau)$, where $\tau=r_p\kappa$ and $\kappa^{-1}$ is the Debye length, was evaluated using the simplified small-Debye-length series approximation when $\tau \geq 3$ [6]. This was the default implementation because all parameter values used in the present study satisfied this condition. The full series representation was also implemented as a numerical option, in which the terms of the infinite series were evaluated explicitly and the remaining integral was computed by adaptive quadrature. The simplified and full calculations gave indistinguishable results over the parameter range used in the present analysis.

*Analytical evaluation of distributed hindrance factors*

Several integrals arising in the distributed pore model were evaluated analytically (see also [1]). In particular, moments of the log-normal pore-size distribution and distributed hydrodynamic hindrance factors were expressed in closed form by approximating the underlying single-pore hindrance functions using Bernstein polynomial expansions.

For a distributed pore system with log-normal pore-radius distribution g(*r*), distributed transport properties can generally be written as

$$\int_a^\infty f(\lambda)\mathrm{g}(r)\mathrm{d}r$$

where *f*(*λ*) represents a hydrodynamic hindrance function and *a* is the solute radius. Direct numerical evaluation of these integrals is computationally expensive and must typically be repeated for each parameter set during nonlinear regression.

To facilitate efficient parameter estimation, the hindrance functions were approximated by Bernstein polynomials,

$$f(\lambda) \approx \sum_{k=0}^{n} \mathrm{c_k B_{k,n}}(\lambda)$$

where $B_{k,n}$ denotes the Bernstein basis polynomial of degree n. Substitution into the distributed transport integrals reduces the problem to weighted sums of incomplete moments of the log-normal distribution, as follows

$$\int_a^\infty f(\lambda)\mathrm{g}(r)\mathrm{d}r \approx \sum_{k=0}^{n}\sum_{j=0}^{n-k} c_k \binom{n}{k}\binom{n-k}{j}(-1)^j a^{k+j} I_{-(k+j)}(a).$$

The incomplete moments of the log-normal distribution are

$$I_m(a) = u^m e^{\frac{m^2 \log(s)^2}{2}} \Phi\left(\frac{\log(u) - \log(a) + m\log(s)^2}{\log(s)}\right)$$

where Φ is the standard normal cumulative distribution function.

The resulting closed-form expressions substantially reduced computational cost while preserving excellent agreement with the original numerical solutions over the physiologically relevant range of pore and solute sizes.

*Hydraulic permeability and large-pore contribution*

The hydraulic permeability parameter $K_f$ was estimated by nonlinear mixed-effects regression. The measured glomerular filtration rate $J_v$, obtained from $^{51}$Cr-EDTA clearance, was used as an input to the model. The effective pressure gradient corresponding to a given $K_f$ was therefore

$$\Delta P = \frac{J_v}{K_f}$$

The fraction of filtrate passing through the large-pore pathway was calculated from the large-pore parameter $a_L$, the total volume flux, and the reflection coefficients of albumin-sized solutes in the small- and large-pore pathways. Albumin was represented by a Stokes–Einstein radius of 3.6 nm and surface charge density of -22 mC/m$^2$.

*Effective pore viscosity*

The viscosity of water was calculated as a function of temperature using the empirical relation

$$\eta(T) = 2.414 \cdot 10^{-5} 10^{247.8/(T-140)},$$

where T is absolute temperature in Kelvin (310°K). To account for confinement effects in narrow pores, the bulk viscosity was multiplied by a steric correction factor,

$$\eta_p = \eta(T) \left( 1 + 18 \left( \frac{r_w}{r_p} \right) - 9 \left( \frac{r_w}{r_p} \right) \right) (1 - f\xi^2),$$

where $r_w$=0.28 nm is the effective radius of a water molecule and $r_p$ is the pore radius. Electrostatic interactions additionally increase the local viscosity through the viscoelectric effect. Following the classical viscoelectric formulation [7], the electric field $\xi$ was estimated from the pore-wall surface charge density using the Grahame equation and the corresponding Debye length. This effective viscosity was used throughout the calculation of hydraulic permeability, diffusion coefficients and Péclet numbers.

*Numerical implementation*

The model was implemented in C and called from R using the .Call interface. Numerical integrations were performed using adaptive Gauss–Kronrod quadrature from the GNU Scientific Library. Free diffusion coefficients were calculated from the Stokes–Einstein relation, and the viscosity of water at 37°C was used with corrections for pore confinement and electroviscous effects.

Nonlinear mixed-effects regression was performed using the *nlme* package in R. Model fitting was performed on log-transformed sieving coefficients. The small-pore radius and geometric standard deviation were fixed as described above. The hydraulic permeability parameter $K_f$ was modeled on the log scale with a random intercept by animal. The electrostatic pore-wall parameter $q_r$ was estimated as a single fixed-effect parameter applied to anionic polysucrose; for neutral polysucrose, $q_s$ = 0, and electrostatic interactions were therefore absent. The large-pore fraction parameter $a_L$ was estimated as a fixed-effect parameter.

Model predictions were compared with observed glomerular sieving coefficients over the measured range of polysucrose radii. Approximate 95% confidence intervals for fixed-effect parameters were obtained from the fitted nonlinear mixed-effects model and back-transformed to the original parameter scale.
